# Accounting for endogenous effects in decision-making with a non-linear diffusion decision model

**DOI:** 10.1101/2022.07.18.498922

**Authors:** Isabelle Hoxha, Sylvain Chevallier, Matteo Ciarchi, Stefan Glasauer, Arnaud Delorme, Michel-Ange Amorim

## Abstract

The Drift-Diffusion Model (DDM) is widely accepted for two-alternative forced-choice decision paradigms thanks to its simple formalism and close fit to behavioral and neurophysiological data. However, this formalism presents strong limitations in capturing inter-trial dynamics at the single-trial level and endogenous influences. We propose a novel model, the non-linear Drift-Diffusion Model (nl-DDM), that addresses these issues by allowing the existence of several trajectories to the decision boundary. We show that the non-linear model performs better than the drift-diffusion model for an equivalent complexity. To give better intuition on the meaning of nl-DDM parameters, we compare the DDM and the nl-DDM through correlation analysis. This paper provides evidence of the functioning of our model as an extension of the DDM. Moreover, we show that the nl-DDM captures time effects better than the DDM. Our model paves the way toward more accurately analyzing across-trial variability for perceptual decisions and accounts for peri-stimulus influences.

## Introduction

Perceptual decision-making has been studied extensively from behavioral^1,2^ and neurophysiological^3,4^ perspectives, as it is omnipresent in daily activities. When decisions are timed, evidence accumulation models accurately describe human and animal behavior. They assume that decisions are made when enough sensory evidence has been gathered.

Among them, the Diffusion Decision Model (DDM, also called Drift-Diffusion Model)^5^ suggests that evidence is accumulated linearly, with a constant drift. The accumulation is additionally subject to Gaussian noise; hence the decision state can be seen as a particle following a Brownian motion. The popularity of this model yields from its intuitive formalism and good fit to behavioral^1^ and neurophysiological data^3^. It has also been shown that the DDM formalizes the optimal strategy for decision-making under time constraints^6,7^. Interestingly, other forms of decision models such as the Leaky-Competing Accumulator model^8^, and attractor models^9,10^ can be formulated equivalently to the DDM under certain constraints^6,11^.

The DDM accounts for global statistics of the behavior by describing the Response Time (RT) distribution and error rate. A major limitation of this model is that it does not consider inter-trial variability. However, behavioral studies have shown that sequential effects^12^ impact prior expectations and the subsequent decision process^13^. Traditionally, expectations are modeled through the starting point, or *bias*, of the accumulation process^5^. Recent accounts have suggested that choice history affects subsequent drift rates^14^. Together, these studies suggest that these parameters could be intertwined and vary over time, as participants become more familiar with the task. To address this issue,^15,16^ proposed an extended DDM, where starting points are uniformly distributed and drifts follow a Gaussian distribution without explicit dependence between them. However, this only provides global statistics about perceptual responses without insight into single decisions or inter-trial interactions. Moreover, the linear accumulation does not describe the variation of the dynamics at the scale of the single decision, which seems inconsistent with the aforementioned empirical observations.

Linear evidence accumulation also assumes that evidence accumulation is independent of the decision state or of the time that passes. While some models consider the effect of time on the decision^17^, or dynamics close to the threshold^18,19^, no model to our knowledge accounts for initial dynamics. For example, ambiguous stimuli could yield flat initial drifts. This is partially translated into non-decision time, as it is assumed to be when sensory evidence is processed in the brain without contributing to the decision process.

Previous attempts at single-trial fitting of decisions have been made through attractor models^9,20,21^, and it has been shown that these models can be reduced to a Drift-Diffusion Model^6,22^, that is in that case, a Langevin equation with a non-linear drift^23^. Its dynamics allow for transitions between decision states under fluctuating stimuli^11^. However, the link between each parameter and the dynamics of the model is complicated to interpret. Moreover, the reduction proposed assumes a reflection symmetry of the network to obtain the given form. This seems limiting, in particular when each perceptual decision recruits different sensory modalities.

Moreover, while the few parameters of the DDM are advantageous in terms of complexity, it can be a limiting factor when analyzing endogenous effects, such as fatigue or training on decisions. Previous works have shown that post-stroke fatigue increases the non-decision time along experiment time, while response times tended to decrease in healthy participants^24^. Increased environmental requirements in terms of workload can decrease RTs and alter accuracy^25^. While some models have taken into account the passing of time within each trials^17^, no models have tried to account for more global fluctuations to our knowledge.

Here we propose a straightforward one-dimensional non-linear form to address these limitations: the non-linear Drift-Diffusion Model (nl-DDM). It recreates double-well-like dynamics from an evidence-accumulation perspective without assuming reflection symmetry. We show its validity and compare its fitting performances to these of the DDM. We first provide a formal description of the nl-DDM, relating it to the DDM. Then, we fit the models on two human behavior datasets: a lexical classification task already published^26^, and a multisensory classification task. Then, we used correlations to compare the parameters of both models on data simulated from DDM parameters and provide an empirical explanation of the effect of the nl-DDM parameters with analogies on the DDM. We show that it fits data equally well as the DDM while providing drift variability like the extended DDM. The dependency of the drift rate on the decision state provides a framework for more refined analyses of the decision process. Last, we considered the time spent performing the lexical task and showed that the nl-DDM modeled behavioral data significantly better than the DDM in that instance, supporting the necessity to account for the experiment time. We provide open-source code pluggable onto the PyDDM toolbox^22^ for reproducibility and easy use of our model.

## Results

In this paper, we introduce the non-linear Drift-Diffusion Model (nl-DDM) and show that it performs better than the DDM in terms of fitting accuracy on behavior. To this aim, we fitted both models on two datasets: a lexical classification dataset previously published in^26^, on which we also modeled the effects of the time spent doing the experiment, and a multi-sensory classification task To provide insight into the meaning of nl-DDM parameters, we also performed correlation analyses between nl-DDM and DDM parameters on data generated from DDM parameters and subsequently fitted by the nl-DDM.

### nl-DDM formalism

Our goal was to propose a simple model in which trajectories are attracted to a boundary. Placing ourselves in the context of two-alternative choice paradigms, our model needed two attractive states. In one dimension, this forces the existence of an unstable fixed-point between the two stable fixed-points^27^.

Therefore, the model we propose follows a Langevin equation where the drift varies with the state of the decision. The drift equation can be written in the following form:

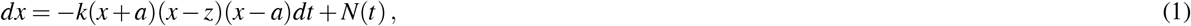

where *x* represents the decision variable and *dx* its variation in infinitesimal time *dt. N*(*t*) is a Gaussian white noise term. −*k*(*x* + *a*)(*x* − *z*)(*x* − *a*) represents the drift, and depends on several parameters. The parameter *k* is a time constant of the system, and *a* and *z* determine where the attractors, or decision boundaries, lie. ±*a* represent the two attractive states, and we constrain *z* to the interval] − *a, a*[to obtain *z* the unstable fixed-point. In this case, the drift corresponds to the deterministic part of the equation and depends on the current decision state. A summary of the parameters of the nl-DDM is given Figure 1, which can be compared to the description of the DDM Figure 5. In the following, we provide a formal explanation of the meaning of each parameter.

**Figure 1.**
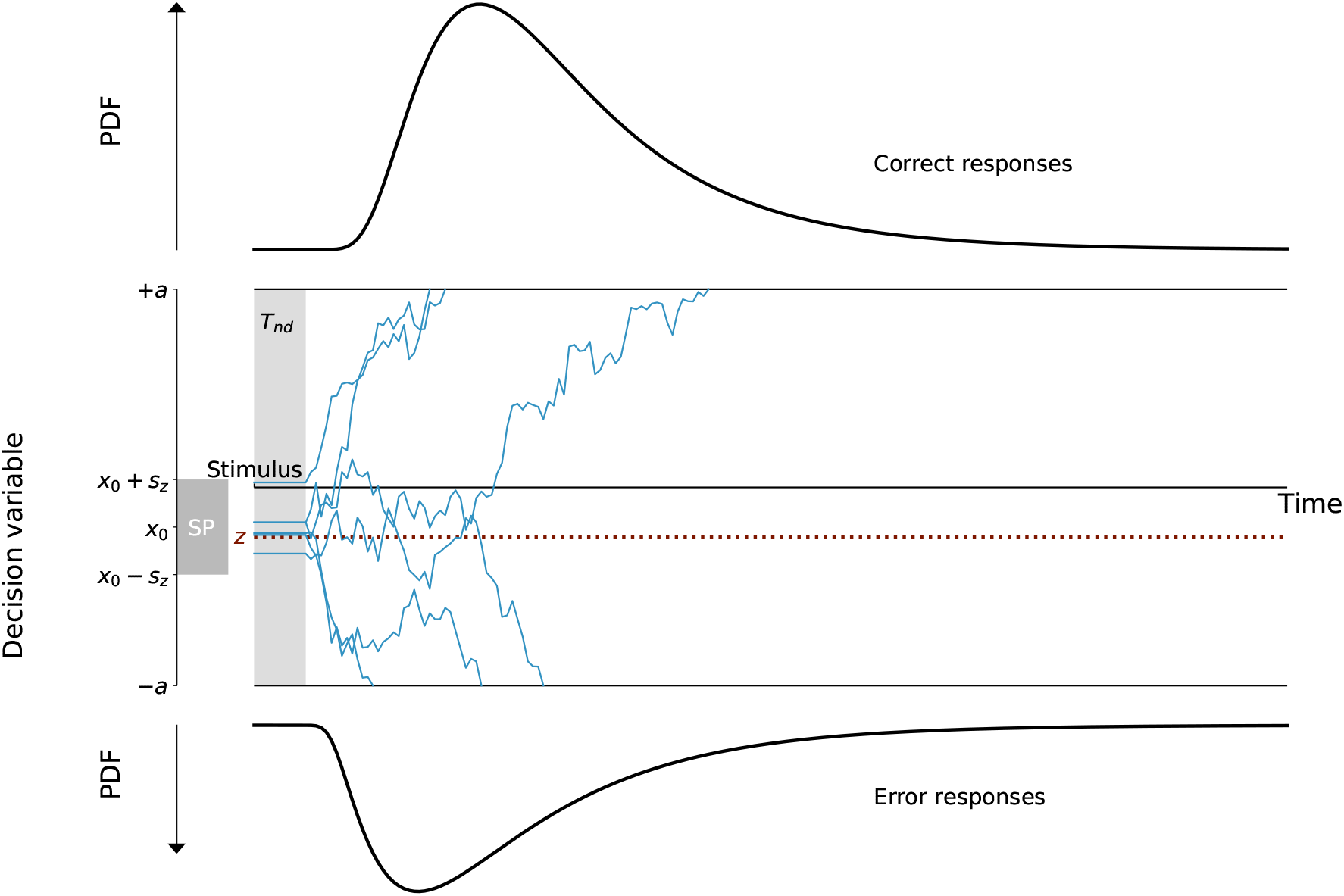
Description of the Non-linear Drift-Diffusion Model (nl-DDM). The decision state is represented by a decision variable *x* traveling from a starting point (for example, drawn from a uniform distribution, centered around *x*_0_ and of width 2*s*_*z*_. It is represented as “SP” on the figure) to a boundary (“Correct boundary” or “Incorrect boundary”) under the influence of a drift. Here, the drift depends on the current state of the decision. Depending on the position of *x*_0_ relative to *z*, the drift will hence have different shapes. The trajectory is also impacted by white noise so that real trajectories are similar to the thin blue lines. From the stimulus onset, the decision process is delayed by a certain non-decision time (*T*_*nd*_). Over an ensemble of decisions, probability density functions of correct and error response times can be created, as displayed here.

The interpretation of *k* as a time constant is straightforward from the equation: as *k* increases, a decision is reached faster for any given set of parameters.

To build an intuition for the other parameters, we first consider the potential function derived from the drift term (Figure 2):

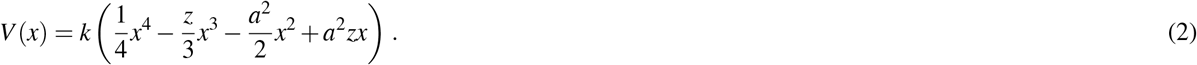

**Figure 2.**
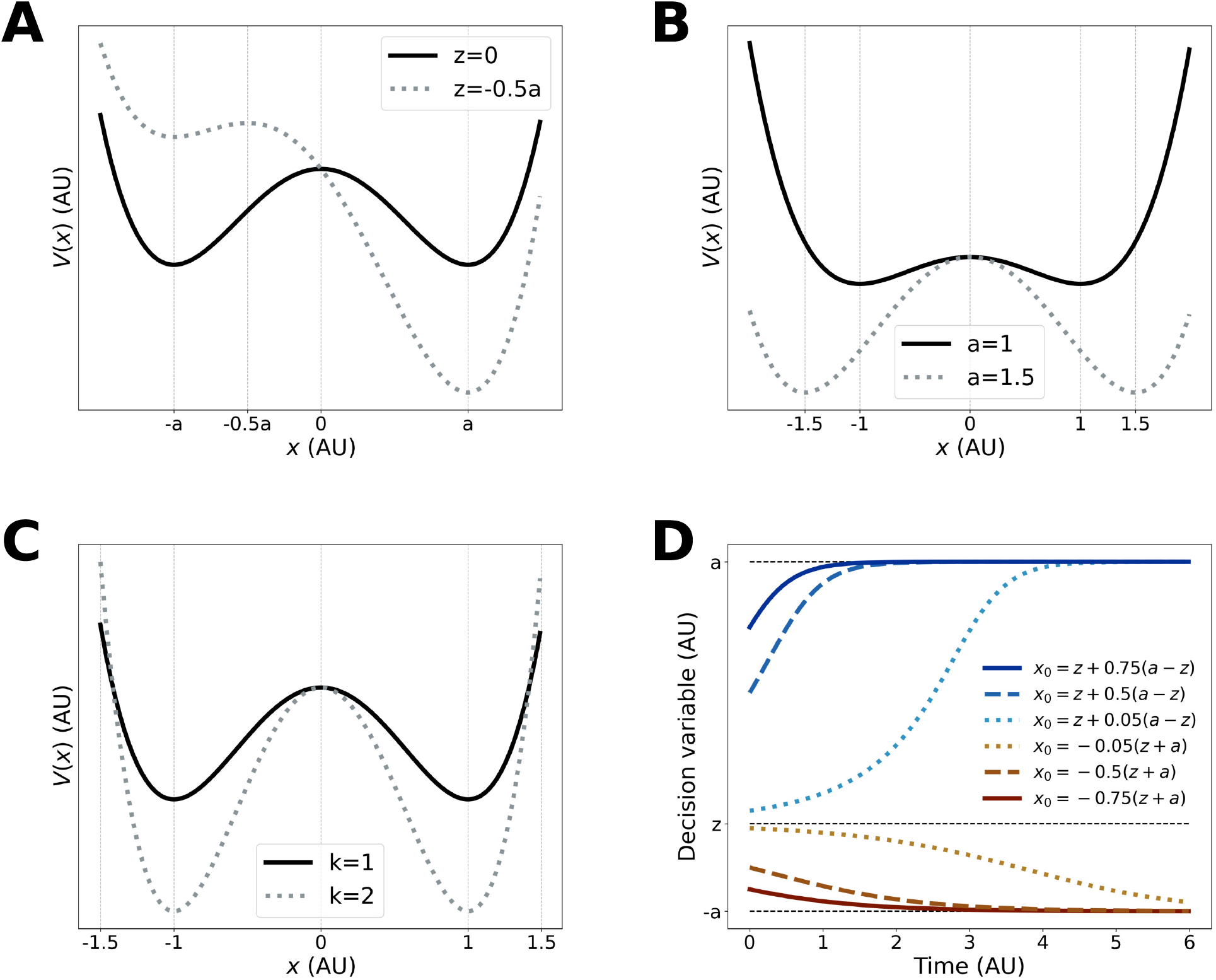
Parameter manipulation on the nl-DDM. A, B, C: Potential functions of the nl-DDM for different *z* (A. Shifting *z* changes the relative attractiveness of each boundary, *a* (B. Shifting *a* changes the accuracy and the speed of decisions), and *k* (C. Shifting *k* changes the speed of decisions). The parameters are always the same for the solid black curve: *a* = 1, *k* = 1, *z* = 0, allowing for a comparison of the effects of the different parameters. D: Trajectories in the absence of noise for different values of *x*_0_, under *a* = 1, *k* = 1, *z* = 0. It becomes clear that the drift range for each trajectory depends on the starting point. The trajectory approaches the boundary asymptotically and will eventually be crossed since noise is omnipresent.

This profile is called a double-well potential profile.

From Figure 2, we can see that there are two sinks at ±*a*, as well as a source at *z*, which emerge from the topology of the system. Therefore, ±*a* are the decision boundaries and control, along with *z*, the speed-accuracy trade-off. Taking *a* the boundary for correct responses and −*a* for incorrect ones, we can see that moving *z* closer to −*a* makes the *a*-well shallower and the −*a*-well deeper (Figure 2A). In other words, the correct decision becomes more attractive than the incorrect one. The gradient becoming more positive on the interval [*z, a*], the trajectories starting on that interval also reach the correct decision faster.

By reducing *a*, both wells become shallower, making decisions slower (Figure 2B). However, for a given noise scale, this also means that any perturbation in the wrong direction is easier to correct because a small perturbation in the other direction can counterbalance that effect. When the wells are deep, the decision variable is then driven rapidly to the stable fixed-point, making perturbations less reversible.

We can also observe the impact of *k* on the potential function in Figure 2C.

Similar to the DDM, we can fit RTs by solving the Fokker-Planck equation corresponding to the Langevin equation (Equation 1)^22^. Then, a non-decision time *T*_*nd*_ shifts the resulting distribution and accounts for biological transmission delays.

This model is similar to the Double-Well Model (DWM), which emerges from attractor network models^11,23^. The potential profile of the DWM indeed takes the form:

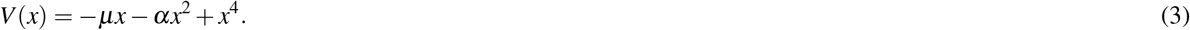

Comparing this equation to Equation (2), we observe a term in *x*^3^ that is absent from the DWM, because of the reflection symmetry assumption made in the DWM^23,27^. However, when *z* = 0 and *μ* = 0, we observe the equivalence of the systems:

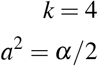

This equivalence is coherent with the interpretation of *z* and *μ* as the impact of the stimulus on the decision and shows that in the absence of a stimulus, the two models follow the same behavior. Because the nl-DDM is not assuming reflection symmetry, the presence of a stimulus impacts the trajectories generated by the two models in different ways.

### Model performance and comparisons

#### Behavioral results

It is helpful to obtain each participant’s RTs and decision accuracy for decision-making analysis, particularly for decision model fitting.

We used two datasets in this paper, described in the Methods section. They both consist of classification tasks performed by human participants. One of them is a dataset collected by Wagenmakers et al. (2008)^26^, in which participants had to assess whether a word presented on screen existed or not. The second one is a dataset in which participants were shown visual stimuli or a combination of visual and auditory stimuli on screen and had to classify them according to their type (either “face” or “number+sound”).

We describe here the validation conducted on the multi-sensory dataset. Analyses of the lexical dataset^26^ are discussed later. On average, participants were shown 49.82±2.42% of “number+sound” stimuli, indicating the quasi-equiprobability of each stimulus. We then performed mixed-model ANOVAs on their RTs and response accuracy for both stimulus-response mapping (between-subject factor) and stimulus (within-subject factor). Across all participants and stimulus types, the mean RT is 535±61 ms (mean ± standard deviation, *N* = 25), with an accuracy of 98.59±0.95%. For the “face” stimulus, participants responded after 539±56 ms with an average accuracy of 98.51±1.17%. Participants responded to the “number + sound” stimulus after 531±69 ms on average with an accuracy of 98.68±0.94%. The difference in performance between the types of stimuli is not significant in terms of accuracy (Table 1) nor RTs (Table 2).

**Table 1.** Within Subjects Effects on Accuracy

| Cases | Sum of Squares | df | Mean Square | F | p |
| --- | --- | --- | --- | --- | --- |
| Stimulus | $1.249 \times 10^{-5}$ | 1 | $1.249 \times 10^{-5}$ | 0.299 | 0.590 |
| Stimulus * S-R mapping | $1.758 \times 10^{-4}$ | 1 | $1.758 \times 10^{-4}$ | 4.202 | 0.052 |
| Residuals | $9.623 \times 10^{-4}$ | 23 | $4.184 \times 10^{-5}$ | | |

**Table 2.** Within Subjects Effects on Response Times

| Cases | Sum of Squares | df | Mean Square | F | p |
| --- | --- | --- | --- | --- | --- |
| Stimulus | 1201.903 | 1 | 1201.903 | 2.446 | 0.132 |
| Stimulus * S-R mapping | 370.446 | 1 | 370.446 | 0.754 | 0.394 |
| Residuals | 11303.230 | 23 | 491.445 |  |  |

In the “face-left” stimulus-response mapping, where participants were instructed to click left upon face stimulus presentation and right when they were presented with a number+sound stimulus, participants responded on average within 531±74 ms with an accuracy of 98.48±1.12% (*N* = 15). Participants (*N* = 10) who underwent the “face-right” mapping responded within 541±30 ms with an accuracy of 98.77±0.60%. The effect of the stimulus-response mapping on accuracy and RT was not significant (Tables 3 and 4). We note a marginal interaction effect between stimulus-response mapping and stimulus type on the accuracy of participants (*p* = 0.052, Table 1).

**Table 3.** Between Subjects Effects on Accuracy

| Cases | Sum of Squares | df | Mean Square | F | p |
| --- | --- | --- | --- | --- | --- |
| S-R mapping | $8.608 \times 10^{-5}$ | 1 | $8.608 \times 10^{-5}$ | 0.447 | 0.510 |
| Residuals | 0.004 | 23 | $1.926 \times 10^{-4}$ | | |

**Table 4.** Between Subjects Effects on Response Times

| Cases | Sum of Squares | df | Mean Square | F | p |
| --- | --- | --- | --- | --- | --- |
| S-R mapping | 1438.081 | 1 | 1438.081 | 0.179 | 0.676 |
| Residuals | 184867.754 | 23 | 8037.728 |  |  |

These results show the uniformity of participant responses across mappings and stimuli.

#### Comparison of loss values

Parameter fitting was performed using PyDDM^22^ for both the nl-DDM and the DDM, minimizing the negative log-likelihood function. As participants in the sensory classification dataset were shown two types of stimuli, we fitted a model per participant and model type, resulting in 25 DDM and 25 nl-DDM fitted. The DDM was fitted using 6 parameters (1 boundary, 2 drifts, i.e. one per stimulus, 1 starting point, 1 starting point variability, 1 non-decision time), and the nl-DDM consisted of 7 parameters (*k, a*, 2*z* (one per stimulus), 1 starting point and variability, 1 non-decision time). We then compared BIC values pairwise.

We computed the Bayesian Information Criterion (BIC) for each model fitted on the multi-sensory dataset to establish a comparison of model performance that takes into account the sample size and number of parameters necessary for each model. This is indeed necessary when comparing performance across model types since the number of parameters is different. We observed that the nl-DDM fitted RT data significantly better than the DDM even when accounting for the number of parameters (Figure 3, Shapiro-Wilk test: *W* = 0.928, *p* = 0.08, one-tailed paired *t*-test, *t*(49) = 1.714, *p* = 0.046, `Cohen’s` *d* = 0.343, *N* = 25).

**Figure 3.**
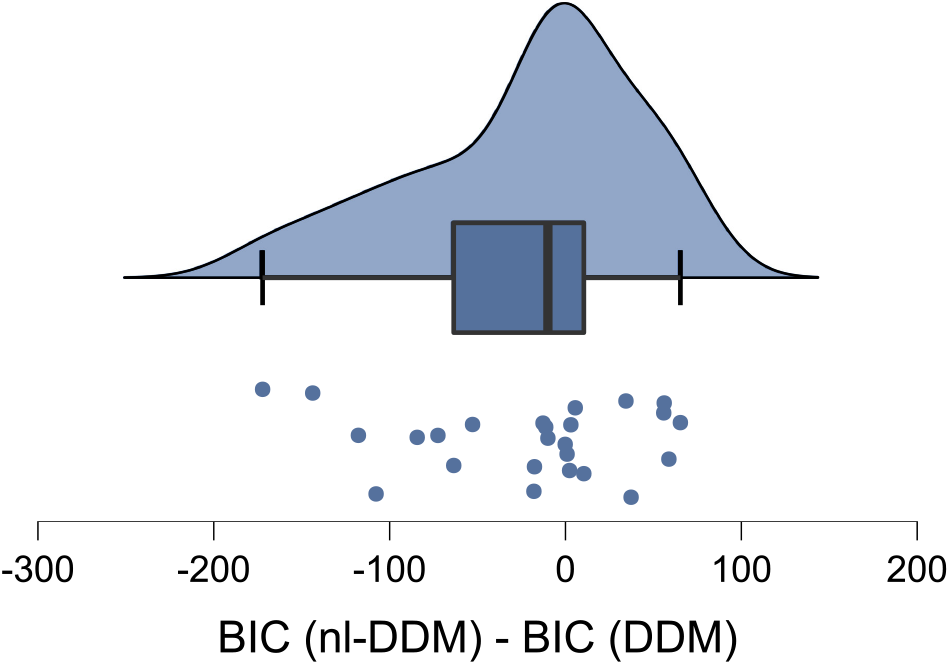
Distribution of the differences between the BIC obtained after fitting the nl-DDM and fitting the DDM on the multi-sensory classification dataset. A more negative difference means a better fit of the nl-DDM compared to the DDM. This figure has been generated using JASP (0.16.0.0)^28^(see https://jasp-stats.org/)

#### Comparison of parameters

We compared the parameters of the DDM and nl-DDM using data generated from DDM parameters, varying the parameters *B, ν, x*_0_ and *s*_*z*_ consecutively. Each parameter varied 100 times, resulting in 400 generated datasets, to which the parameters *k, z, a, x*_0_ and *s*_*z*_ of the nl-DDM were fitted.

The correlation matrix of the nl-DDM and DDM parameters across all models is given Figure 4. Note that, since the DDM parameters were artificially varied, the correlation coefficients within DDM parameters were discarded from our analysis.

**Figure 4.**
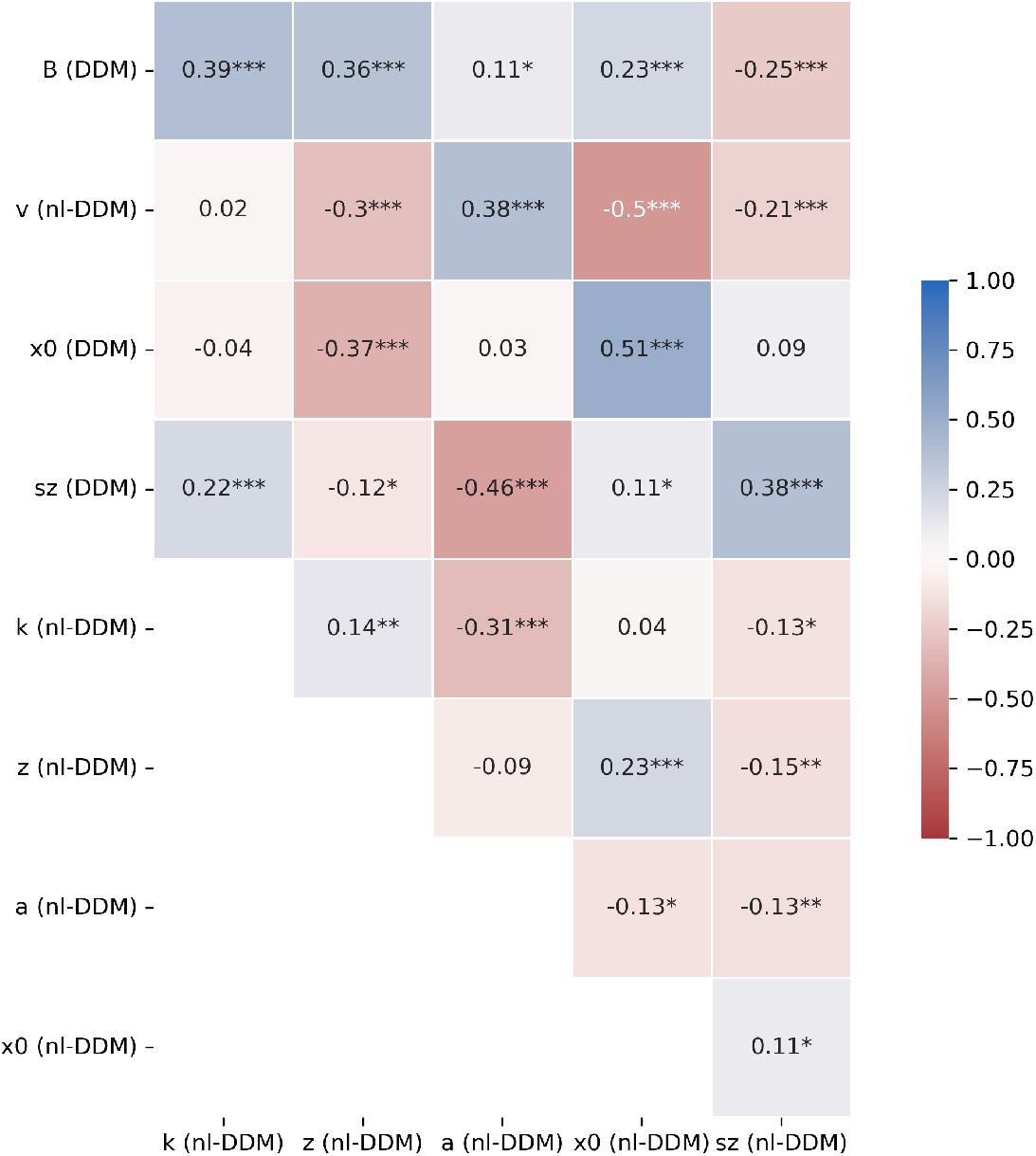
Correlation matrix of all parameters, computed from the parameters fitted over the multi-sensory classification dataset. Pearson correlation coefficients were computed over *N* = 50observations. This figure was obtained using the matplotlib (3.5.2)^30^-based Python library seaborn (0.11.2)^31^ (see https://matplotlib.org/ and https://seaborn.pydata.org/). ^⋆^ : *p* < 0.05,^⋆⋆^ : *p* < 0.01,^⋆⋆⋆^ : *p* < 0.001.

We empirically compute the relation between the parameters within the nl-DDM. We observe a strong negative correlation between *a* the boundary and *k* the time constant. This corresponds to their opposed effect on the attractiveness of the correct response. Increasing either will make the decision more attractive, so to keep the same attractiveness of the correct response, if one increases, the other should decrease. In our data, since the noise term is kept constant, these two terms are strongly correlated. Note that the effect of each parameter is still different, as shown on Figure 2B and C. While increasing *k* deepens both wells, increasing *a* will deepen the wells and pull them apart. Effectively, the relation between *a* and *k* is not linear (see Supplementary Figure S2). We note a positive correlation between *k* and *z*, which can also be understood in reference to Figure 2A and C: if *k* increases, the correct decision well becomes deeper and thus more attractive, and to correct for this effect, *z* needs to increase as well. *k* and *s*_*z*_ the width of the starting point distribution of the nl-DDM are related, while *a* relates both to the middle of the distribution *x*_0_ and to it width *s*_*z*_. This can be explained by the symmetric effect of *k* on the depth of the potential wells (Figure 2C) and the asymmetric effect of *a* (Figure 2B). In the same line, *z* correlates positively to *x*_0_ and negatively to *s*_*z*_. Increasing *z* results in slower correct and less accurate decisions. To maintain the same speed-accuracy trade-off, the starting point distribution can be shifted towards the correct decision boundary and it variability diminished so that a larger portion of it is located to the right of *z*, which is the attracting zone of the correct decision boundary. Last, the starting point parameters *x*_0_ and *s*_*z*_ in the nl-DDM are positively correlated with each other.

Upon cross-model comparison, we first observe that the middles of the starting point distributions and their width are positively correlated, which was expected. *x*_0_ of the nl-DDM and *s*_*z*_ of the DDM are consequently also positively correlated, since they both correlate positively to *s*_*z*_ of the nl-DDM, and *a* negatively to *s*_*z*_ of the DDM for the same reason. *x*_0_ of the DDM also correlates negatively with *z* of the nl-DDM: increasing *x*_0_ in the DDM results in faster correct decisions and more accurate decisions. Decreasing *z* in the nl-DDM has the same effect, as the correct decision will becomes deeper and a larger proportion of the starting point distribution is located to the right of *z*, hence to the side of the correct decision. *s*_*z*_ of the DDM additionally correlates negatively to *z* and positively to *k*. Increasing *s*_*z*_ while having *x*_0_ = 0 in the DDM results in faster decisions and to a diminished accuracy. Decreasing *z* in the nl-DDM result in faster correct decisions, which is coherent with the effect of increasing *s*_*z*_. Increasing *k* makes decisions faster and since the potential wells become deeper (Figure 2C), the decisions are also more prone to noise and hence less accurate. All these effect mirror the ones observed upon increasing *s*_*z*_ in the DDM.

The DDM boundary *B* is correlated positively to *k, a, z* and *x*_0_ and negatively to *s*_*z*_ of the nl-DDM. Increasing the DDM boundary results in an improved accuracy at the cost of slower decisions. Consistent with this effect, increasing *a* in the nl-DDM also results in more accurate decisions. Shifting the starting point distribution towards the correct decision, i.e. to the right of *z*, while decreasing its variability results in more correct decisions (Figure 2D), explaining the observed correlation. and increase in *z* results in slower correct responses due to the correct potential well being shallower (Figure 2A), which mirrors the loss of speed implemented by an increase of the decision boundary in the DDM. Surprisingly, an increase in *k* results in overall faster and less accurate decisions, which contradicts the effects of the boundary increase in the DDM. However, we also noted that *k* correlated positively with *z* and negatively with *s*_*z*_ of the nl-DDM. These two parameters being positively and negatively correlated to *B* respectively, the increase of *k* upon increasing of *B* should be a consequence of these correlations.

We also observe a significant negative correlation between *z* and *ν*. This relationship was also expected, as increasing the drift *ν* in the DDM results in faster correct decisions. Mirroring this effect, *z* regulates the relative attractiveness of each decision well. As *z* becomes more negative, the correct decision (corresponding to decision boundary +*a*) becomes more attractive, and hence correct decisions are made faster. Another explanation for this can be derived from Figure 2: if we shift *z* closer to 0, the negative and positive wells of 2 will tend to be at the same level. It means that the mean maximum drift will decrease towards zero as *z* increases closer to the middle of the two boundaries ±*a*. In other words, increasing *z* will decrease the drift, hence the negative correlation. Conversely, *a* correlates positively with *ν* since pulling the potential well apart makes them more attractive (Figure 2B). *z* also correlates positively with *x*_0_ of the DDM, which is expected as both have an impact on the proportion of trials that reach either boundary in the absence of noise. Last, we note a negative correlation between DDM drift and nl-DDM starting point distribution. Following^29^, in the DDM framework higher starting point variability results in faster error responses. Therefore, across models, if the drift becomes greater in the DDM, hence making error responses slower, the starting point variability in the nl-DDM should diminish to have the same effect.

### Effects of time passing

The time spent performing the task is likely to impact the decision strategy, and we seek to model these effects using the DDM and the nl-DDM. We thus added a time condition to the lexical classification dataset, corresponding to whether the trial was performed in the first half of the experiment (“early” condition) or the second (“late” condition). One of the 17 participants did not complete all the blocks (participant 2), and was eliminated from our analyses. Therefore, the following analyses are presented over 16 subjects, each exposed to all the conditions.

#### Model fitting

The drift and *z* of the DDM and nl-DDM varied as a function of the stimulus complexity (common, rare, very rare, non-existent word), and the boundary of the DDM varied depending on both the instruction (speed or accuracy) and the time condition, resulting in 4 boundaries. In the nl-DDM the effects were modeled separately using *a* and *k*, fitting *k* depending on the time condition and *a* according to the instruction. Then, the starting point distribution and non-decision time were fitted over all trials for each model. We compared the fitting performance of the two models. The BIC of the nl-DDM was significantly smaller than the BIC obtained by the DDM (Shapiro-Wilk test: *W* = 0.773, *p* < 0.001, one-sided Wilcoxon signed-rank test (*BIC*_*nl*−*DDM*_ < *BIC*_*DDM*_: *W* = 34, *p* = 0.042), which shows that splitting the effects of instruction and experiment time yielded better results than combining them.

### Analysis of the parameters of the lexical classification dataset

We fitted both the DDM and the nl-DDM taking into account the instruction, the time of the experiment (early or late trial) and the word type for each trial. In the DDM, we hence fitted 4 drifts, corresponding to the 4 word types, and 4 boundaries, corresponding to `2instructions ×2times`. Conversely in the nl-DDM, we fitted 4*z* parameters (one per word type), 2*a* (one per instruction) and 2*k* (one per time of the experiment). We then performed paired *t*-tests to assess the discriminability of the parameters across conditions. We first compared the drift and *z* parameters across word types. In both the DDM and nl-DDM, we observed significant differences in the drifts and *z* between all word type pairs, except between rare and non-existent words (Tables 5,6). The two models therefore discriminate between word types equally well. We observed that *k* differed significantly between the two time conditions (*t*(15) = 4.553, *p* < 0.001), and *a* differed significantly between the two instruction conditions (*t*(15) = 4.879, *p* < 0.001). Comparing the boundaries of the DDM resulted in 6 comparisons, corresponding to the Bonferroni-corrected 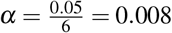. We noted no significant difference between early and late trials in the accuracy instruction (*t*(15) = 2.784, *p* = 0.014), while all the other differences were significant (Table 7). The behavioral analysis (see Supplementary Information 3) however underlined significant differences in response times and accuracy across time of the experiment. The effects of time passing are therefore better transcribed by the nl-DDM than the DDM.

**Table 5.** Paired Samples T-Test on the values of *z* fitted per word type on the lexical classification dataset

| Measure 1 |  | Measure 2 | t | df | p | Cohen's d |
| --- | --- | --- | --- | --- | --- | --- |
| $z_{\text{frequent}}$ | - | $z_{\text{rare}}$ | -7.526 | 15 | < .001 | -1.882 |
| | - | $z_{\text{very rare}}$ | -7.173 | 15 | < .001 | -1.793 |
| | - | $z_{\text{non-existent}}$ | -5.438 | 15 | < .001 | -1.360 |
| $z_{\text{rare}}$ | - | $z_{\text{very rare}}$ | -3.933 | 15 | 0.001 | -0.983 |
| | - | $z_{\text{non-existent}}$ | 1.082 | 15 | 0.296 | 0.271 |
| $z_{\text{very rare}}$ | - | $z_{\text{non-existent}}$ | 5.617 | 15 | < .001 | 1.404 |

**Table 6.** Paired Samples T-Test on drift values fitted per word type on the lexical classification dataset

| Measure 1 |  | Measure 2 | t | df | p |
| --- | --- | --- | --- | --- | --- |
| $v_{\text{frequent}}$ | - | $v_{\text{rare}}$ | 7.257 | 15 | < .001 |
| | - | $v_{\text{very rare}}$ | 15.695 | 15 | < .001 |
| | - | $v_{\text{non-existent}}$ | 9.899 | 15 | < .001 |
| $v_{\text{rare}}$ | - | $v_{\text{very rare}}$ | 16.337 | 15 | < .001 |
| | - | $v_{\text{non-existent}}$ | 0.419 | 15 | 0.681 |
| $v_{\text{very rare}}$ | - | $v_{\text{non-existent}}$ | -10.031 | 15 | < .001 |

**Table 7.** Paired Samples T-Test on the decision boundary of the DDM, fitted on the lexical classification dataset according to the instruction (BA*i*: boundaries for the accuracy instruction, BS*i*: boundaries for the speed instruction) and time of the experiment (B*x*1: early trials, B*x*2: late trials)

| Measure 1 |  | Measure 2 | t | df | p | Cohen's d |
| --- | --- | --- | --- | --- | --- | --- |
| BA1 | - | BA2 | 2.784 | 15 | 0.014 | 0.696 |
|  | - | BS1 | 9.635 | 15 | < .001 | 2.409 |
|  | - | BS2 | 10.884 | 15 | < .001 | 2.721 |
| BA2 | - | BS1 | 8.003 | 15 | < .001 | 2.001 |
|  | - | BS2 | 10.295 | 15 | < .001 | 2.574 |
| BS1 | - | BS2 | 7.020 | 15 | < .001 | 1.755 |

## Discussion

We presented a non-linear model of decision-making. This model takes the form of a Langevin equation, and provides a framework in which individual trajectories of the decision variable can have different shapes under the same global parameters (Figure 2D). We have shown that this model predicts behavioral data equally well as the DDM. From the formalism we have described, it becomes clear that inter-trial variability in drift emerges from the dynamics of the nl-DDM, offering the possibility for further single-trial analyses and modeling.

The interpretation of the nl-DDM parameters may seem counter-intuitive at first, particularly when considering that decisions are made faster when the boundaries are further apart. Our correlation analysis provided insight into bridging the meaning of nl-DDM and DDM parameters. The difference is that in the DDM, the gradient of the drift is constant, whereas it varies in decision space with the nl-DDM. By pulling the boundaries further apart, we effectively reduce the impact of one attractor on the other, making each of them more attractive. Therefore, a decision can be reached faster, at the price of accuracy. Similarly, increasing the drift in the DDM is equivalent to shifting *z* towards the negative boundary in the nl-DDM, as they both result in fast correct responses. However, note that these parameters are not entirely equivalent as we did not find a perfect mapping between them, meaning that the nl-DDM is conceptually different from the DDM.

We have shown that the nl-DDM can also account for changes in the decision dynamics that do not relate directly to a change in the experiment but rather to the state of the participants. While such an account necessitates an exponential multiplication of parameters in the DDM, the nl-DDM requires a simple duplication of the parameters. Indeed, if we again consider the case where there are two conditions (here, instruction and time), the DDM requires *n*_instructions_ ×*n*_times_ boundaries, while the nl-DDM requires *n*_instructions_ + *n*_times_ boundary parameters (*a* and *k*). The version we showed here only split the trials into two instruction and time conditions, which results in the same number of parameters in the DDM and in the nl-DDM. Note however that any more instances of either condition would have meant more parameters in the DDM relative to the nl-DDM. We argue that drift and starting point variability are intertwined, as transcribed in the nl-DDM and in alignment with the view that evidence accumulation starts in anticipation of stimulus apparition^32^. EEG research has shown a matching between pre-stimulus activity and confidence ratings in human participants^33,34^. The starting point distribution models pre-stimulus states, and in the DDM the drift relates to the quality of the integrated stimulus^4^, with more ambiguous stimuli corresponding to lower drift rates. At the single-trial level, drift variability relates to the variation of the quality of stimulus perception and processing in the brain^1^. In our model, the starting point directly impacts evidence accumulation, allowing for a uniform theory of decision-making that includes explicit co-dependency of parameters. Some general forms of the DDM include a variance of the drift. In the nl-DDM, we have not implemented this possibility, as we assumed that the inter-trial variability of the drift emerged from the variability of the starting point. In neurophysiological terms, we assumed that the pre-stimulus arousal and stimulus expectations led to differences in the rate of evidence accumulation. This is supported by past observations, according to which pre-stimulus brain activation impact RTs^35,36^. Pre-stimulus brain activity also modifies perceptual^37^ and pain^38^ thresholds. Therefore, depending on the pre-stimulus activity, decisions can be made even in the absence of actual evidence^33,39^, or under ambiguous evidence^40,41^. Along the same lines,^42^ have shown that biases were implemented through local changes in accumulation rate, which supports the intertwining of accumulation rate and pre-stimulus states. However,^34,43–45^ argue that pre-stimulus brain states should only affect the decision criterion, not how well participants perceived the stimuli. Translating the signal-detection theory to the evidence-accumulation scheme^16^, pre-stimulus states should only be changing the decision boundary, or equivalently the starting point, and not the drift rate. For example,^34^ found that pre-stimulus alpha power did not impact the accuracy of visual evidence accumulation, but only the confidence in the decision.^33^ found similar results with auditory stimuli. Although these observations seem to contradict our assumption that the starting point impacts the evidence-accumulation process, both phenomena could co-exist, as more extreme starting points are more attracted to the closer attractor. This results in fast and confident observations, although little evidence has been accumulated (we would be located at a plateau in our model), i.e., even if the stimulus was not well perceived.

The current analysis uses a form of the DDM without all variabilities proposed by Ratcliff and Tuerlinckx^29^. The reason is two-fold. First, we wanted to use simple forms of both models to emphasize the characteristics of the ground parameters of each model. One may argue that the DDM should then have been fitted using a single point as the starting point. However, this would have introduced a confound when comparing the two models. Indeed, a major advantage of the nl-DDM is the variety of dynamics that it offers depending on the starting point. Since the DDM can also be improved by adding starting point variability, implementing it in both models seemed to be a fair compromise. Second, and related to the first, any source of variability that could have been introduced in the DDM could also have been implemented in the nl-DDM. Besides starting point variability, non-decision time variability could also exist in the nl-DDM. As argued above, the drift and starting point variability should be intertwined, meaning that fitting a variability of the drift would be redundant to some extent in the nl-DDM, but we could implement variability in *k*, which, according to our analyses on time that passes, could be a natural mirror of the remaining effects of drift variability in the DDM.

The dynamics that we propose here is rooted in empirical observations made in neurophysiological studies. More specifically, three phases can be identified in the decision trajectories: an inertia stage, a quasi-linear evidence accumulation stage, and a plateau stage. The initial inertia relates directly to the brain activation needed to integrate sensory evidence.^35^ and^36^ have shown in human EEG studies that the brain activity prior to stimulus presentation changed the speed of responses. More specifically, they showed that the more pre-activated the required sensory area, the faster the decision. The nl-DDM mimics this behavior at the single-trial level: for trials starting close to the unstable fixed-point (i.e., further from the correct decision well), the trajectories start with a plateau-like stage, whereby little evidence is accumulated because the brain would need to process the stimulus more intensively in order to extract information from it, before integrating evidence faster. This initial inertia is circumvented by shifting the starting point closer to the decision well, resulting in faster and more accurate responses. The initial inertia in the DDM is referred to as the non-decision time and encompasses both sensory processing and motor planning and execution. The nl-DDM assumes therefore that part of these processes participates in the decision process, which goes beyond the conceptualization of decision-making as the sequence of sensation, perception, and motion.

A recent review from^46^ shows the limitations of existing evidence-accumulation models. We try to address several of them with the nl-DDM, including the possibility for analyses beyond the global description of RTs and the formulation of initial and final dynamic changes during the decision process. In particular, our formal description has shown that different shapes of decision trajectories can co-exist within the same framework, not solely because of noise, but because of meaningful variability. We expect this model to be further used to gain insight into the across-trial variability of decisions.

The current study considered that the input was presented at the beginning of the trial and affected the decision in a constant fashion. We could also imagine more dynamic cases, where the input is processed over a finite period and participants accumulate evidence during stimulus presentation, as has been done in past DDM analyses^47,48^. In non-stationary contexts, the input can be considered as a variation of *z* in time. By shifting *z* to either boundary, more trajectories are attracted to the opposite boundary, hence increasing the likelihood of correct answers. In addition, it can be inferred from our formal analysis that changing *z* means changing the drift rate. This change in input could also explain error-correcting behaviors^49^ and spontaneous changes of mind^50^. When the stimulus ends, the DDM is modified so that the drift is null, i.e. evidence is no longer accumulated. Therefore, changes of mind are the result of noise in the system. Conversely, stimulus termination could be modeled through shifting *z* in the nl-DDM, which effectively modifies the drift rate of the current decision, in a way that the decision variable could toggle towards the opposite boundary upon stimulus disappearance. Conceptually, the drift in the nl-DDM not only relates to the accumulation of evidence but also encompasses decision processes related to the post-processing of evidence.

We have shown that, while similar to the DWM^11^ derived from attractor models^23^, the nl-DDM is equivalent to it only in the absence of input. A question that remains open is that of the mechanism underlying this equation. From the reduction computed in the paper by^23^, it would seem that a network of three populations could produce the dynamics we have described. However, the main assumption of the reduction was that the network was invariant through reflection. We argue that the mechanisms described by the nl-DDM are similar to these of the DWM, but offer a broader range of applications beyond the case of symmetrical models.

Extending this model to multiple-choice situations is another interesting ground of research. The DDM is inapplicable in such situations. The nl-DDM would require structural changes for multiple choices. Indeed, the decision variable’s trajectory is here modeled in a one-dimensional space, where the alternatives are represented as attractors. Its multiple-choice variant would require more attractors. In 1*D*-space, adding more stable fixed-points will result in two issues. First, traveling from one alternative to another may require passing through other decision wells, which seems incoherent with behavior. Second, adding stable fixed-points requires the implementation of as many unstable fixed-points, which would mean a two-fold increase in the number of parameters when adding one choice. A simpler solution would be to switch to a 2*D*-space, so there could be a central unstable fixed-point, and the subjective preference for each alternative would determine the position of each stable fixed-point.

## Methods

### Drift-Diffusion Model

The Drift-diffusion model^5^ is characterized by a linear accumulation disturbed by additive noise. Formally, this can be written as the following Langevin equation (Equation (4)):

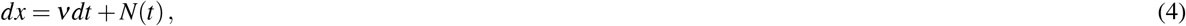

where *x* represents the decision variable, an abstract quantity representing the state of the decision, *dx* its infinitesimal variation in time *dt*, and *N*(*t*) is a Gaussian white noise, parameterized by its standard deviation *σ*. Figure 5 gives a representation of this model.

**Figure 5.**
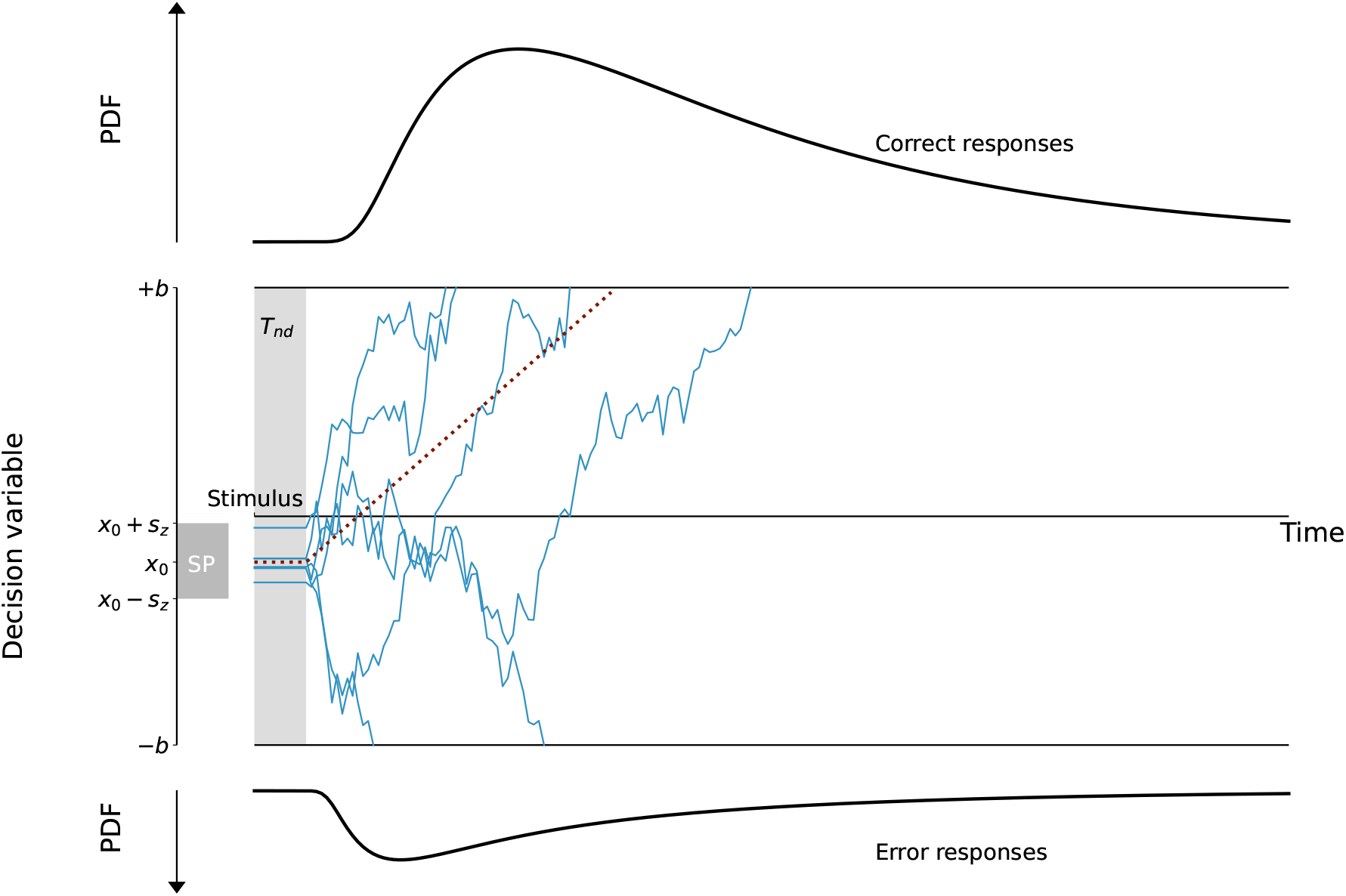
Description of the Drift-Diffusion Model (DDM). The decision state is represented through a decision variable that travels from a starting point that can be drawn for example from a uniform distribution, centered around *x*_0_ and of width 2*s*_*z*_. The decision state is represented through a decision variable *x* traveling from a starting point (for example, drawn from a uniform distribution, centered around *x*_0_ and of width 2*s*_*z*_. It is represented as “SP” on the figure) to a boundary (“Correct boundary” or “Incorrect boundary”) under the influence of a constant drift (dotted line). The trajectory is also impacted by white noise so that real trajectories are similar to the thin blue lines. From the stimulus onset, the decision process is delayed by a certain non-decision time (*T*_*nd*_). Over an ensemble of decisions, RT distributions of correct and error responses can be estimated, as displayed here.

Evidence is accumulated following Equation (4) until a decision boundary *A* > 0 or −*A* is reached. Typically, the positive boundary corresponds to correct decisions and the negative one to incorrect responses.

Finally, the starting point of accumulation is called the bias and is defined as a single point within the two boundaries. In general forms of this model, it is also possible to consider that the starting point is drawn from a uniform distribution centered around the bias *x*_0_ and of width 2*s*_*z*_, such that [*x*_0_ −*s*_*z*_, *x*_0_ + *s*_*z*_] ⊆] −*A, A*[^51^, or from other parametric distributions^15^. We will consider uniformly distributed starting points in our fitting to provide a fair comparison of the two models without loss of generality.

The boundary separation represents the speed-accuracy trade-off. Indeed, if this separation is bigger, decisions are less impacted by noise and hence more accurate, but at the same time, they will take longer to reach from a given starting point. In contrast, the drift mainly impacts the speed of response, as a higher drift will lead to faster correct responses and longer incorrect responses.

Fitting is typically done globally over RTs. In fact, the trajectories defined by the equation cross the decision boundaries, forming a RT distribution usually compared to an exponentially modified Gaussian. In order to obtain a close fit, it is necessary to define a non-decision time (noted *T*_*nd*_), which corresponds to the time necessary for sensory processing of the stimulus, motor planning and execution, independently of the decision process.

### Data collection and processing

In order to test the quality of the fitting of the nl-DDM, we use RTs from a classification task performed by humans described thereafter. The paradigm was initially implemented to assess the relation between RTs and emotion valence of visual stimuli.

#### Classification task with different sensory modalities

We first tested the quality of the nl-DDM by fitting it to data we collected. 25 (11 female, 14 male) healthy right-handed participants aged 27.72 ±8.96 (mean ±standard deviation) with normal or corrected-to-normal vision and hearing consented to taking part in a perceptual classification task experiment. EEG brain activity was also recorded (not reported here). The experiment was performed under the local ethics committee approval of the Comité d’Ethique de la Recherche Paris-Saclay (CER-Paris-Saclay, invoice notice nb. 102). All the methods described were performed in accordance with the guidelines and regulations stated by this committee and disclosed in the invoice. An interview preceded the experiment to check with the participants for non-inclusion criteria (existing neurological and psychiatric disorders, uncorrected visual and hearing deficiencies). Informed consent was obtained from all the participants included in this study.

Participants were presented at each trial with images of faces or images of numbers, and had to respond with mouse clicks to report what stimulus they perceived. A sound accompanied images of numbers to suppress any ambiguity. Participants were instructed to respond using their right hand. To control for possible differences in motor response speeds between the two fingers, one group of participants (*N* = 15) was instructed to report faces with a left click and numbers with a right click (“face-left” stimulus-response mapping), while the other (*N* = 10) was given the opposite instruction (“face-right” stimulus-response mapping). Responses were constrained to two seconds after stimulus onset. No feedback on the performance was given to participants. At each trial, each stimulus had a 50% chance of occurring.

Each participant performed 480 classification trials, split into 8 blocks of 60 trials each. Between each block, participants were offered a break of free duration. Each trial followed the sequence described Figure 6. First, a central red cross appeared on the screen, indicating a pause period. After 1.5 second, the cross became white as a signal for trial start. The white cross stayed for 1.5 second, after which a video clip of visual noise appeared: 9 frames of noise of 100 ms each were displayed. After the noise clip, a last frame of random visual noise was presented, and the stimulus appeared on top of it. The last frame stayed intact until the end of the trial, and the stimulus was displayed over it for 200*ms*. The trial was terminated upon participant response or timed out after 2 seconds. A trial lasted for about 5 seconds, resulting in blocks of about 5 minutes each.

**Figure 6.**
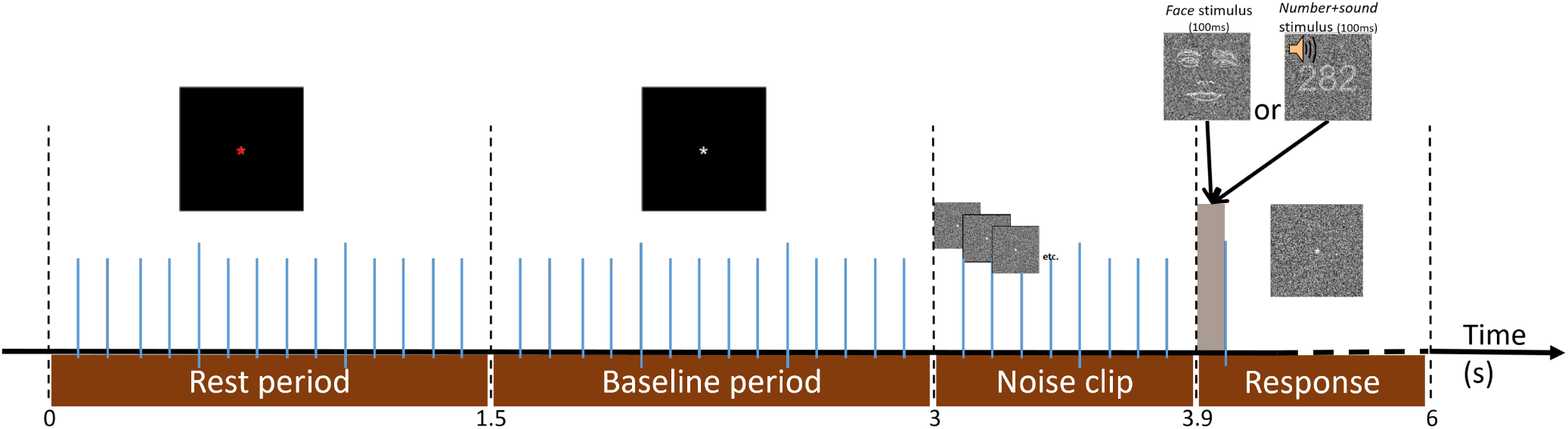
Timeline of a single trial. Each trial is preceded by a rest period, followed by a baseline period (necessary for EEG processing, not reported here), each lasting 1.5 seconds. A noise clip consisting of 9 random-dot frames of 100 ms each indicates the arrival of the stimulus in a non-stimulus-specific fashion. The stimulus then appears on a noisy visual background for 100 ms. The same noisy background frame then lasts until the participant’s response and times out after 2 seconds otherwise.

We used face sketches as used in^52^, which were generated from the Radboud Face Dataset^53^. Number stimuli were generated at the beginning of the session for each participant, under the constraint that they were 3-digit integers. In total, 10 different face stimuli and 10 different number stimuli were used for each participant.

#### Lexical classification dataset from Wagenmakers et al. (2008)^26^

To model the effects of time passing and discard the possibility of better performances emerging from the fitting algorithm or data acquisition, we also lead our analyses on a bigger pre-existing dataset taken from^26^. 17 human participants performed a classification task, as they were randomly presented with real or invented words. The invented words were generated from real words by changing a vowel, and the real words were labeled in three categories depending on their frequency (frequent, rare, or very rare). In total, stimuli were split into 4 categories of interest. Each participant performed 20 blocks of 96 trials each, with as many invented words as real words in each block. Participants were given the additional instruction to define the speed-accuracy trade-off in each block: they alternated between blocks where speed was emphasized and blocks where accuracy was more important. Responses were limited to 3 seconds, and trials with RTs below 180 ms were discarded to avoid anticipatory responses. More details can be found in^26^, and the dataset can be accessed from here.

### Behavioral analyses

We are interested in comparing model parameters between the DDM and the nl-DDM. It is important to check whether participants’ performance across stimulus-response mappings and stimuli is coherent in terms of RTs and accuracy. Indeed, the multi-sensory experimental paradigm we defined entails two types of stimuli and two motor commands for the choices. In addition, we have created two experimental groups, which were instructed to respond with opposite motor commands. First, we computed the percentage of stimuli in each class to verify that the stimuli were globally equiprobable for each participant. Since we designed the experiment to display each stimulus with the same probability at each trial, we expect this number to be close to 50%. Otherwise, participants could opt for a strategy that prioritizes one response against the other. Then, we performed two mixed-model ANOVAs, respectively testing RTs and accuracy. The stimulus-response mapping was considered a between-subject factor and the stimulus type a within-subject factor.

In the lexical classification data^26^, the effect of the time of the experiment is of special importance, as well as its interaction with the other experimental conditions (i.e. the word type and the instruction). We therefore performed two repeated-measures ANOVAs, respectively testing RTs and mean accuracy, assessing the effects of time (first half of the trials or second, resulting in two conditions: early vs. late trials), stimulus frequency, and instruction (accuracy or speed). One of the participants (participant 2) did not perform the 9^*th*^ block of the experiment, which removed a significant portion of trials in one of the conditions. This participant was removed from the analyses, and the analyses were therefore performed over 16 participants. Post-hoc analyses were performed using the Holm correction.

### Data fitting

The classical way of fitting evidence-accumulation models is by fitting one drift for each stimulus category separately. In that case, the positive and negative boundaries still correspond to correct and incorrect responses respectively, and the starting points are taken from the same distribution regardless of the stimulus. Consequently, one pair of boundaries ±*B*, the middle of the starting point distribution *x*_0_ and its half-width *s*_*z*_, and two drifts *ν*_0_ and *ν*_1_ (corresponding respectively to “face” and “number+sound” trials) have to be fitted in the DDM. Similarly, one pair of stable fixed-points ±*a*, one time scale *k*, the middle of the starting point distribution *x*_0_ and its half-width *s*_*z*_, and two unstable fixed-points *z*_0_ and *z*_1_ (that will tune the drift in the “face” and “number+sound” stimuli respectively) are needed for the nl-DDM. In both cases we fix the noise parameter to *σ* = 0.3. As explained by^5^, since the speed-accuracy trade-off is determined by the boundary separation, fitting two parameters among drift, boundary, and noise is constraining enough. In addition, each model requires fitting a non-decision time *T*_*nd*_ per stimulus type. Hence, 6 parameters must be fitted per participant for the DDM, against 7 for the nl-DDM.

We used the PyDDM toolbox^22^ for the fitting, minimizing the negative log-likelihood function and an implicit resolution. The nl-DDM indeed does not have explicit solutions when *z* is not centered. The log-likelihood is such that the more negative, the closer the modeled distribution of RTs is to the empirical RT histogram.

#### Fitting the lexical classification dataset^26^

With this dataset, we were interested in modeling the effects of time passing throughout the experiment. Therefore, we created an artificial condition based on the sequence of blocks, which characterized the trials as happening early (i.e. within the first half of the experiment) or late (i.e. within the second half of the experiment). We discarded participant 2, for whom we did not have data for the 9^*th*^ block. All the other participants performed 20 blocks alternating between speed and accuracy instructions, therefore each time condition held 5 blocks with each instruction. Within the DDM framework and for each participant, one drift was computed per stimulus type, resulting in 4 drift terms: *ν*_1_, *ν*_2_, *ν*_3_, *ν*_*NW*_, corresponding respectively to frequent, rare, very rare, and non-existent word stimuli. 4 boundaries were fitted, corresponding to 2 instructions ×2 time conditions. The non-decision time, starting point, and starting point variability were fitted for each participant over all trials. The within-trial noise parameter was fixed to 0.3. Hence, each model consisted of 11 parameters.

Within the nl-DDM framework, we fitted for each participant one *z* per stimulus type (*z*_1_, *z*_2_, *z*_3_, *z*_*NW*_). In addition, one parameter *a* was fitted per instruction condition and one parameter *k* per time condition, resulting in 4 more parameters. Similar to the DDM, the non-decision time, starting point, and starting point variability were fitted over all trials from each participant and the noise scale is set to 0.3.

As previously, we used PyDDM^22^ with negative log-likelihood minimization and implicit resolution.

#### Performance comparison

Since the fitting on both datasets was performed using a different number of parameters and samples, we computed the Bayesian Information Criterion for each model, defined as:

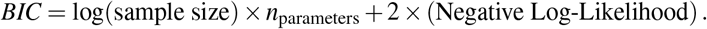

That way, a penalty for more samples and parameters is considered. The negative log-likelihood is the fitting score.

Hence, we compare each loss pairwise, using a one-sided paired-sample *t*-test. Indeed, we want to test whether the nl-DDM is better than the DDM with these three metrics, hence testing the hypothesis BIC_nl-DDM_ < BIC_DDM_.

### Comparison of parameters

For a better understanding of the parameters of the nl-DDM, their interaction and their meaning in the DDM framework, we computed the Pearson’s correlation coefficients of DDM and nl-DDM parameters over simulated experiments. In order to obtain correlation coefficients within nl-DDM parameters as well as across DDM and nl-DDM parameters, we varied one by one the DDM parameters *B, ν, x*_0_ and *s*_*z*_, and simulated 500 data points for each parameter combination. The nl-DDM parameters *k, z, a, x*_0_ and *s*_*z*_ were subsequently fit to the generated datasets. Table 8 summarizes the sampling of DDM parameters as well as the default value for each parameter. We explored 100 variations of each parameter, resulting in 400 generated datasets. Since the noise parameters *σ* = 0.3 and *T*_*nd*_ = 0.3*s* were kept constant when simulating the DDM, we also fixed *σ* = 0.3 and *T*_*nd*_ = 0.3 in the nl-DDM.

**Table 8.** List of DDM parameters used for simulations and subsequent correlation analysis. A uniform sampling of 100 values was performed over the variation range of each parameter. One parameter was varied at a time, using the default value for all the other parameters.

| Parameter | Default value | Variation range |
| --- | --- | --- |
| B | 1 | [0.2, 5] |
| $v$ | 0.2 | [0, 10] |
| $x_0$ | 0 | [-1, 1] |
| $s_z$ | 0 | [0, 1] |
| $\sigma$ | 0.3 | Kept constant |
| $T_nd$ | 0.3 | Kept constant |

We computed the correlation matrix between all the parameters of both models. This allows for a first look into first-order interactions between model parameters, within and across model types. Since the correlations within DDM parameters were irrelevant due to their artificial manipulation, these coefficients were not computed. The correlation coefficients were computed using Pearson’s *ρ*, defined as:

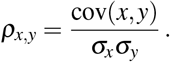

## Acknowledgements

The authors thank Nicolas Nieto for his discussions on the fitting method.

## Author contributions statement

IH: conception of the model; data analysis; interpretation; drafting; revisions. SC: conception of the model; data analysis; revisions. MC: conception of the model; revisions. SG: conception of the model; revisions. AD: revisions. MAA:data acquisition; revisions.

## Additional information

### Data availability

The lexical classification dataset is already made available by Wagenmakers et al. (2008)^26^ and can be accessed here. The multi-sensory classification dataset as well as the Python code will be made available upon acceptance of this work on a public repository. The corresponding author can be contacted regarding these datasets.

