## Supplementary Information for "Accounting for endogenous effects in decision-making with a non-linear diffusion decision model"

### 1 Comparing the Ornstein-Uhlenbeck and the nl-DDM model

Other non-linear models of decision-making exist, among which the Ornstein-Uhlenbeck (OU) model[1]. This model takes the form:

$$dx(t) = (\lambda x(t) + \mu)dt + N(t)$$

We note that, similar to the nl-DDM, the variation depends on the current state of the decision $x(t)$. $\mu$ corresponds to the effect of the stimulus, identical to the effect of the drift in the DDM. $\lambda$ represents the effect of the state of the participant on the accumulation process. To represent an accumulation process for a two-alternative forced-choice task, accumulation boundaries have to be fitted in addition to these parameters. To compare its fitting performance to these of the nl-DDM, we fitted both models on the multi-sensory classification dataset. The nl-DDM was fitted in the same way as presented in the main paper: one pair of stable fixed-points $\pm a$, one time scale $k$, the middle of the starting point distribution $x_0$ and its half-width $s_z$, and two unstable fixed-points $z_0$ and $z_1$ (that will tune the drift in the "face" and "number+sound" stimuli respectively), a non-decision time, and the noise scale was set to $\sigma = 0.3$. We fitted $\lambda$, $\mu_0$ and $\mu_1$ (corresponding to the two sensory stimuli), a boundary $B$, a non-decision time, the middle of the starting point distribution $x_0$ and its half-width $s_z$, and the noise scale was also set to $\sigma = 0.3$. In total, $x$ parameters were fitted for each model. The fitting performances were assessed using the Negative Log-Likelihood (since all models are fitted using the same number of parameters and the same number of samples, the BIC is simply a shifted version of the Negative Log-Likelihood). We observe that the nl-DDM offers a slightly better description of data, although not significant (Negative Log-Likelihood(nl-DDM)-Negative Log-Likelihood(OU)= $-1.096 \pm 21.206$, mean±standard deviation, paired-sample $t$-test: $t(49) = 0.258, p = 0.798$).

### 2 Correlation analysis on empirical data

The goal of the correlation analysis presented in the main paper was to underline links between the DDM and nl-DDM parameters. In this part, we are interested in knowing whether these relations are also found empirically.

We computed the Pearson's correlation coefficients of the nl-DDM parameters over all conditions and participants, using only the multi-sensory classification dataset for simplicity, i.e., over $N = 50$ observations. This allows supporting the observations we have noted in the formalism part. Indeed, since fewer parameters were fitted in this case than for the lexical classification dataset, the comparison becomes more straightforward. From the 25 participants, we obtained 50 fits per model type by duplicating for each stimulus type the boundaries and time constant terms, hence separating the stimulus types and obtaining $25 \times 2$ fits per model type.

The results of this analysis are presented Figure S1 and S2. The correlations within nl-DDM parameters observed on simulated data are found again. We observe that some of the correlations across nl-DDM and DDM parameters are modified due to DDM parameters not being fixed, as opposed to the analysis on simulated data presented in the main article.

### 3 Behavioral analysis of the lexical classification dataset[4]

The tables of ANOVAs on the behavior held by participants on the lexical classification task are given Tables S1, S2 and S3. We assessed the effects of word type, instruction and time of the experiment on both RT and accuracy by performing repeated-measures ANOVAs.

The RT varied significantly with word type ($F(3,45) = 36.329, p < 0.001$, Supplementary Table S1), instruction ($F(1,15) = 16.541, p = 0.001$), and time ($F(1,15) = 55.260, p < 0.001$), with a significant interaction effect of word type and instruction ($F(3,45) = 3.305, p = 0.021$), word type and time ($F(3,45) = 82.976, p < 0.001$), and word type, time and instruction ($F(3,45) = 9.579, p < 0.001$). Post-hoc comparisons revealed that non-existent words resulted in significantly shorter RTs than any other word type, frequent words shorter RTs than rare or very rare words, and rare words yield prompter responses than very rare words (Supplementary Table S3).As expected, participants also responded significantly faster in the speed condition compared to the accuracy condition ($t(15) = 4.067, p_{Holm} = 0.001$). Interestingly, participants responded faster in late relative to early trials ($t(15) = 7.434, p_{Holm} < 0.001$).

The accuracy was also significantly impacted by word type ($F(3,45) = 179.581, p < 0.001$, Supplementary Table S2), instruction ($F(1,15) = 23.863, p < 0.001$), time ($F(1,15) = 102.297, p < 0.001$), and the interaction between word type and time ($F(3,45) = 54.412, p < 0.001$), and word type, time and instruction ($F(3,45) = 3.163, p = 0.034$). Post-hoc analyses revealed that both frequent and non-existent words were responded to more accurately than both rare and very rare words (frequent-rare: $t(15) = 18.567, p_{Holm} < 0.001$, frequent-very rare: $t(15) = 11.197, p_{Holm} < 0.001$, non-existent-rare: $t(15) = 19.874, p_{Holm} < 0.001$, non-existent-very rare: $t(15) = 12.504, p_{Holm} < 0.001$). Participants also responded significantly more accurately to rare words compared to very rare words ($t(15) = 7.370, p_{Holm} < 0.001$). Accuracy trials were more accurate on average than speed trials ($t(15) = 4.885, p_{Holm} < 0.001$). Early trials were more accurate than late trials ($t(15) = 10.114, p_{Holm} < 0.001$).

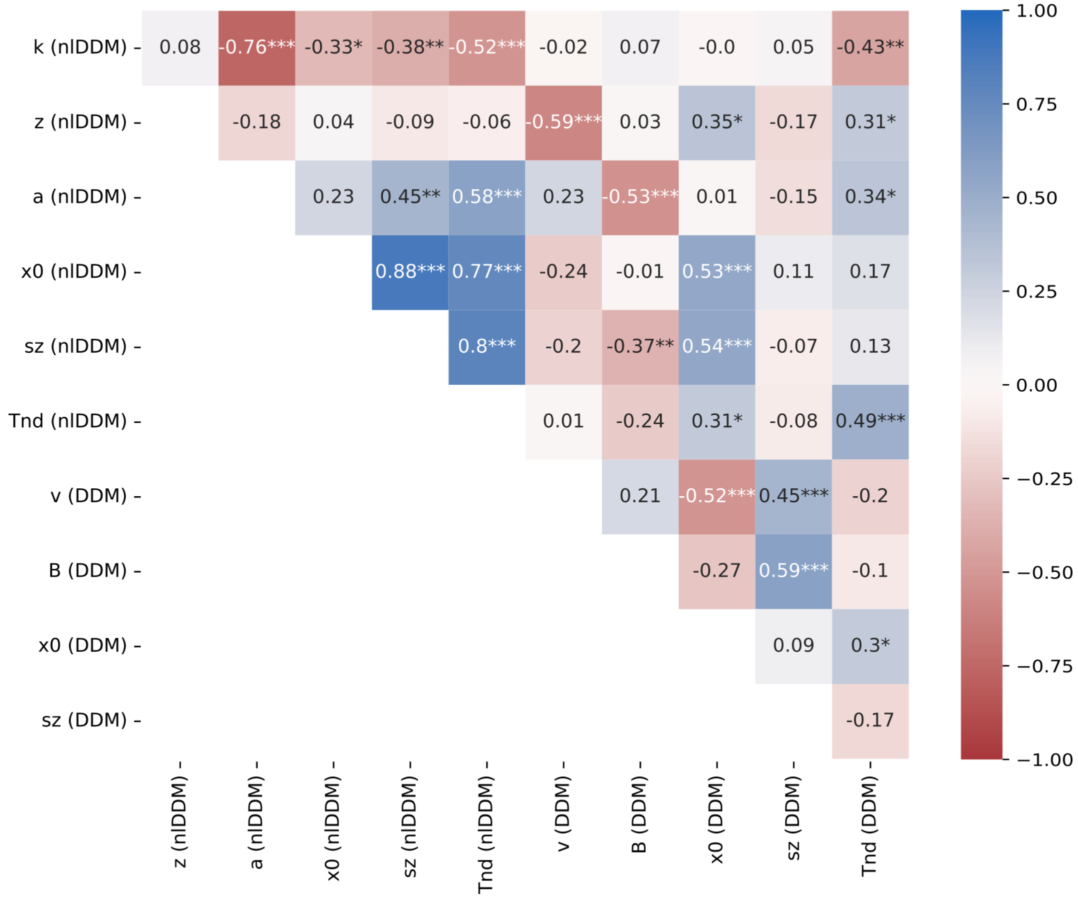


**Figure S1.** Pearson correlation coefficients between DDM and nl-DDM parameters, nl-DDM and DDM parameters fitted over the multi-sensory dataset. This figure was obtained using the matplotlib (3.5.2)[2]-based Python library seaborn (0.11.2)[3] (see https://matplotlib.org/ and https://seaborn.pydata.org

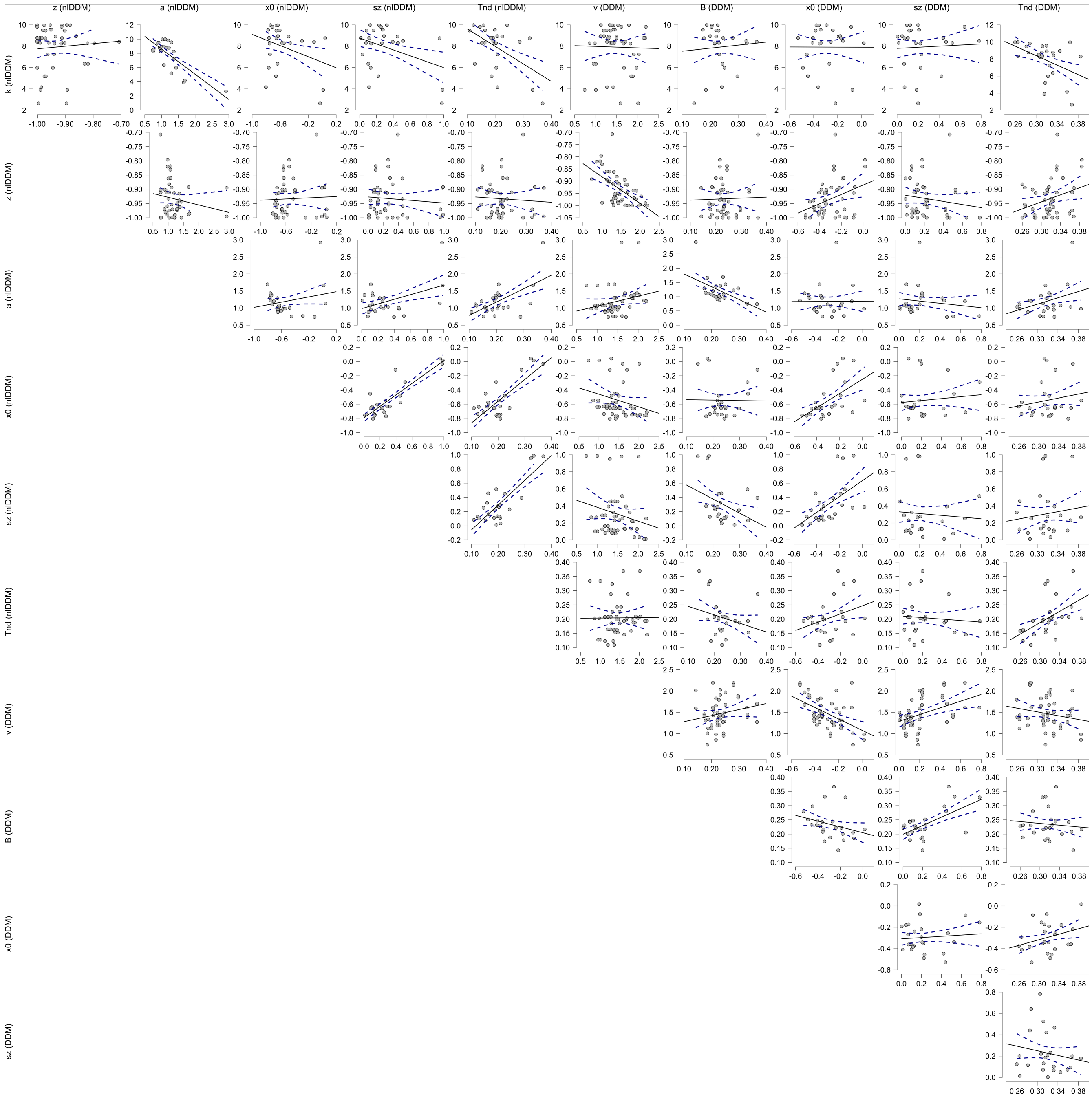


**Figure S2.** Correlation plots of DDM and nl-DDM parameters, computer from the fitting over the multi-sensory classification dataset. The correlations were computed as Pearson's correlation coefficient.The blue dashed lines represent the 95% confidence intervals for the regression. This figure has been generated using JASP (0.16.0.0)**?**(see https://jasp-stats.org/). [NOTE: This figure has been added since the previous version]

From these results, it is noteworthy that the time of the experiment significantly impacts behavior, and seems to imply modifications in the speed-accuracy trade-off strategy. Note also that there is no interaction effect between time and instruction.

**Table S1.** ANOVA on Response Times of the lexical classification task

| Cases | Sum of Squares | df | Mean Square | F | p |
|---|---|---|---|---|---|
| Word type | 2.998 | 3 | 0.999 | 36.329 | $< .001$ |
| Residuals | 1.238 | 45 | 0.028 | | |
| Time | 0.073 | 1 | 0.073 | 55.260 | $< .001$ |
| Residuals | 0.020 | 15 | 0.001 | | |
| Instruction | 0.044 | 1 | 0.044 | 16.541 | 0.001 |
| Residuals | 0.040 | 15 | 0.003 | | |
| Word type * Time | 0.427 | 3 | 0.142 | 82.976 | $< .001$ |
| Residuals | 0.077 | 45 | 0.002 | | |
| Word type * Instruction | 0.021 | 3 | 0.007 | 3.223 | 0.031 |
| Residuals | 0.099 | 45 | 0.002 | | |
| Time * Instruction | $7.895e-4$ | 1 | $7.895e-4$ | 0.882 | 0.363 |
| Residuals | 0.013 | 15 | $8.950e-4$ | | |
| Word type * Time * Instruction | 0.023 | 3 | 0.008 | 9.579 | $< .001$ |
| Residuals | 0.035 | 45 | $7.843e-4$ | | |

**Table S2.** ANOVA on Mean accuracy of the lexical classification task

| Cases | Sum of Squares | df | Mean Square | F | p |
|---|---|---|---|---|---|
| Word type | 6.591 | 3 | 2.197 | 179.581 | $< .001$ |
| Residuals | 0.551 | 45 | 0.012 | | |
| Time | 0.057 | 1 | 0.057 | 102.297 | $< .001$ |
| Residuals | 0.008 | 15 | $5.549e-4$ | | |
| Instruction | 0.108 | 1 | 0.108 | 23.863 | $< .001$ |
| Residuals | 0.068 | 15 | 0.005 | | |
| Word type * Time | 0.230 | 3 | 0.077 | 54.412 | $< .001$ |
| Residuals | 0.063 | 45 | 0.001 | | |
| Word type * Instruction | 0.001 | 3 | $4.380e-4$ | 0.337 | 0.799 |
| Residuals | 0.059 | 45 | 0.001 | | |
| Time * Instruction | 0.001 | 1 | 0.001 | 2.195 | 0.159 |
| Residuals | 0.010 | 15 | $6.618e-4$ | | |
| Word type * Time * Instruction | 0.008 | 3 | 0.003 | 3.163 | 0.034 |
| Residuals | 0.039 | 45 | $8.717e-4$ | | |

**Table S3.** Post Hoc Comparisons of Word type effect on response times in the lexical classification dataset

| | | Mean Difference | SE | t | $p_{holm}$ |
|---|---|---|---|---|---|
| non-existent | frequent | −0.061 | 0.029 | −2.097 | 0.042 |
| | rare | −0.276 | 0.029 | −9.410 | $< .001$ |
| | very rare | −0.193 | 0.029 | −6.587 | $< .001$ |
| frequent | rare | −0.214 | 0.029 | −7.314 | $< .001$ |
| | very rare | −0.132 | 0.029 | −4.491 | $< .001$ |
| rare | very rare | 0.083 | 0.029 | 2.823 | 0.014 |
